# PhaLP 2.0: extending the community-oriented phage lysin database with a SUBLYME pipeline for metagenomic discovery

**DOI:** 10.64898/2025.12.08.692814

**Authors:** Alexandre Boulay, Victor Németh, Bjorn Criel, Michiel Stock, Bernard De Baets, Clovis Galiez, Elsa Rousseau, Yves Briers, Roberto Vázquez

**Author notes:** These authors contributed equally (Alexandre Boulay, Victor Németh). Instituto de Productos Lácteos de Asturias (IPLA-CSIC), Oviedo, Asturias, Spain.

## Abstract

As biology becomes increasingly data-driven, so too does the field of phage lysins, enzymes that degrade bacterial cell walls and hold promise as alternatives to traditional antibiotics. Five years ago, we introduced PhaLP, a centralized resource for **Pha**ge **L**ytic **P**rotein sequences and associated metadata to support global research efforts. Here, we present PhaLP 2.0, a significantly enhanced database designed to overcome key challenges in the computational study of lysins by integrating newly identified lysins obtained from thousands of metagenomes.

To expand the known diversity of lysins beyond those from cultured phages, we developed SUBLYME, a protein embedding-based machine learning **S**oftware designed to **U**ncover and classify **B**acteriophage **Ly**sins in **Me**tagenomic datasets. Using embeddings derived from the prior well-curated protein sequences of the original PhaLP database, we trained support vector machines to distinguish lysins from non-lysins in viromes and classify them as either endolysins or virion-associated lysins. The models achieved an average F1-score of 98% on held-out lysin clusters. SUBLYME enabled the discovery of 743,000 new lysin sequences from EnVhogDB, a virome-derived protein database, increasing the number of known lysin clusters by a factor of 40, from 1,000 to 40,000. PhaLP 2.0 entries were annotated by integrating Pfam functional predictions to the refined delineations obtained with SPAED, an algorithm that leverages the predicted aligned error matrix from AlphaFold predictions to identify domain boundaries.

Both SUBLYME and the PhaLP 2.0 database are accessible online at https://github.com/Rousseau-Team/sublyme and http://phalp.ugent.be, respectively. Together, these advances establish PhaLP 2.0 as a comprehensive and scalable portal for the discovery, classification, and sequence analysis of phage lysins, paving the way for future antibacterial applications and evolutionary insights.

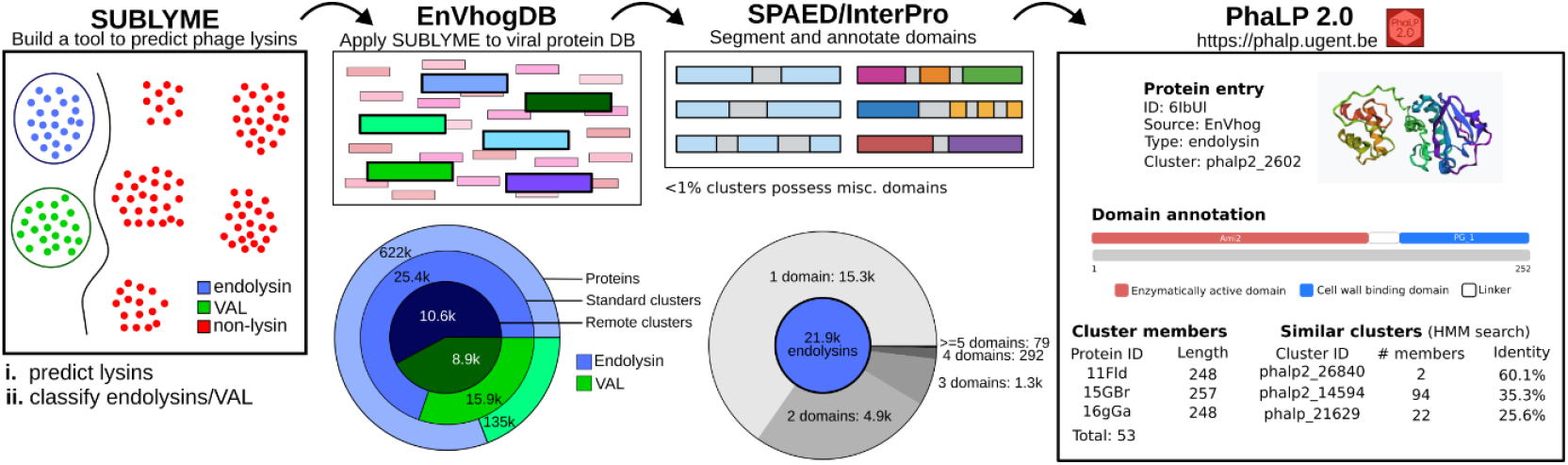

## Introduction

Bacteriophages, or phages, are viruses that specifically infect bacterial hosts (Yutin 2018; Dion 2020). They exhibit remarkable genetic and structural diversity and play a vital role in driving bacterial evolution and shaping microbial ecosystems. Among the proteins encoded by double-stranded DNA (dsDNA) phages, lysins are of particular interest due to both their biological roles and therapeutic potential. These enzymes that degrade the bacterial cell wall may act at two distinct stages of the phage infection cycle: virion-associated lysins (VALs) facilitate genome injection during adsorption, while endolysins mediate host cell lysis at the end of the cycle (Danis-Wlodarczyk 2021; Abdelkader 2022; Bałdysz 2024).

Endolysins, especially those found in Gram-positive infecting phages, often exhibit a modular organization (Fischetti 2010; Oechslin 2022). They are usually composed of two types of functional modules: enzymatically active domains (EADs) and cell-wall-binding domains (CBDs). Modular endolysins are composed of at least one EAD and a CBD, but they can possess multiple modules of each type. For their part, globular endolysins, consisting of a single EAD, are typical of phages that infect Gram-negative hosts (Oliveira 2013). VALs feature more complex and larger architectures, and besides the mandatory EAD (and, on occasions, a CBD) they incorporate structural modules intended for the assembly of the enzyme as part of the viral particle (Criel 2021).

The prevailing view is that the functional properties of endolysins, typically defined in terms of lytic activity and substrate specificity, are largely determined by their modular composition (Schmelcher 2012; Vázquez 2021; Oechslin 2022). The recombination of modules not only enhances their evolutionary adaptability but also enables endolysins to acquire new functions or modify existing ones (Smug 2023). This also allows for the combinatorial engineering of variants with desired characteristics as novel antibacterials, for example, to synthesize endolysins that target a specific bacterial species or to optimize their lytic activity (Cheng 2007; Diez-Martinez 2015; Oechslin 2022). The screening of combinatorial libraries of increasing complexity has already been undertaken in a manner analogous to hit-to-lead development processes, with the aim of identifying lysins that exhibit desirable drug-like properties. In response to the globally increasing interest in lysin research, the PhaLP database was introduced in 2021 as a comprehensive and centralized database of phage lysins (Criel 2021). PhaLP was envisioned to serve as a central resource for lysin research in a data-extensive era and was composed mainly of lysins derived from cultured phages and curated entries from UniProt (UniProt Consortium 2025). However, to capture a more complete view of the diversity of lysins, it is essential to incorporate metagenomic data, which better represents phages infecting unculturable hosts. The EnVhog database is the largest database of phage proteins to date, collected from thousands of environmental metagenomic datasets (Pérez-Bucio 2025). By organizing proteins into groups of remote homologs, EnVhog provides an ideal resource to identify a vast diversity of lysins.

Traditional function annotation methods are often inadequate for metagenomic datasets (Pérez-Bucio 2025; Boulay 2025b) because they contain a much larger diversity than that of current curated databases, thereby limiting the sensitivity of homology-based approaches. An alternative approach that has been the focus of recent studies makes use of protein embeddings (Flamholz 2024; Boulay 2025b). Protein embeddings are vectorial representations of protein sequences obtained from protein language models such as ProtT5 (Elnaggar 2022). These models have been trained on extensive protein sequence corpora and have been shown to capture information on protein functions well. As they are not dependent on the alignment of sequences, embedding-based methods are often much more sensitive for protein classification (Yang 2018; Malec 2025).

Here, we present SUBLYME, a machine-learning-based **S**oftware for **U**ncovering **B**acteriophage **Ly**sins in **Me**tagenomic datasets. Using protein embeddings of the well-curated lysins in PhaLP 1.0 we trained two support vector machines (SVMs) to first identify lysins and subsequently classify them as either VALs or endolysins. We then applied SUBLYME to the EnVhog database, resulting in the creation of PhaLP 2.0, a metagenomic extension of the original resource.

PhaLP 2.0 is thus an enriched database comprising 754k dereplicated lysin entries from both UniProt (11k already in PhaLP 1.0) and EnVhog (743k new entries), complemented by user-friendly tools for advanced querying. It supports customizable searches based on a wide array of filters and attributes, and provides extensive sequence, physicochemical, and structural information for each entry. To further enhance the utility of PhaLP 2.0, we applied SPAED, our previously developed algorithm for 3D structure-based domain delineation (Boulay 2025a) to obtain accurate boundaries of the functional modules within each PhaLP 2.0 entry. We anticipate that PhaLP 2.0, enriched by SUBLYME, will serve as a valuable, community-oriented resource for advancing lysin research, enabling the exploration of previously inaccessible diversity and supporting the development of lysin-based therapeutics in the fight against antibiotic resistance.

## Methods and materials

### Training SUBLYME models

The training dataset for SUBLYME was compiled from two sources (**Figure 1A**). First, lysins, including both endolysins and VALs, were obtained from the well-curated PhaLP database (v2023_01, accessed 2023/08/07) (Criel 2021). Second, non-lysins were obtained as described in Boulay et al. (Boulay 2025b) from GenBank using INPHARED (Cook 2021). The protein-coding genes present in the phage genomes acquired with INPHARED (accessed 2024/01/02) were predicted using Prodigal (v2.6.3) (Hyatt 2010) and annotated using PHROGs (Terzian 2021). Proteins with no annotations or annotations related to lysins, such as peptidase, amidase, lysozyme and deacetylase, were removed (see complete list in the create_dataset.ipynb notebook in our Zenodo archive). All proteins were clustered at 30% sequence identity and 70% coverage with MMseqs2 (version 17-b804f) (Steinegger 2017). These parameters are the standard for identifying homologous proteins (Pearson 2013). Only one randomly selected representative protein per cluster was kept for non-lysins, finally obtaining 15,387 lysins (1,100 clusters) and 47,288 non-lysins (i.e., from 47,288 clusters) that form the dataset used to train and test SUBLYME. ProtT5 (prot_t5_xl_half_uniref50-enc) (Elnaggar 2022) was used to compute the embeddings of all these proteins. The protein sequences, embeddings, functional annotations and cluster information are available in our Zenodo archive.

**Figure 1.**
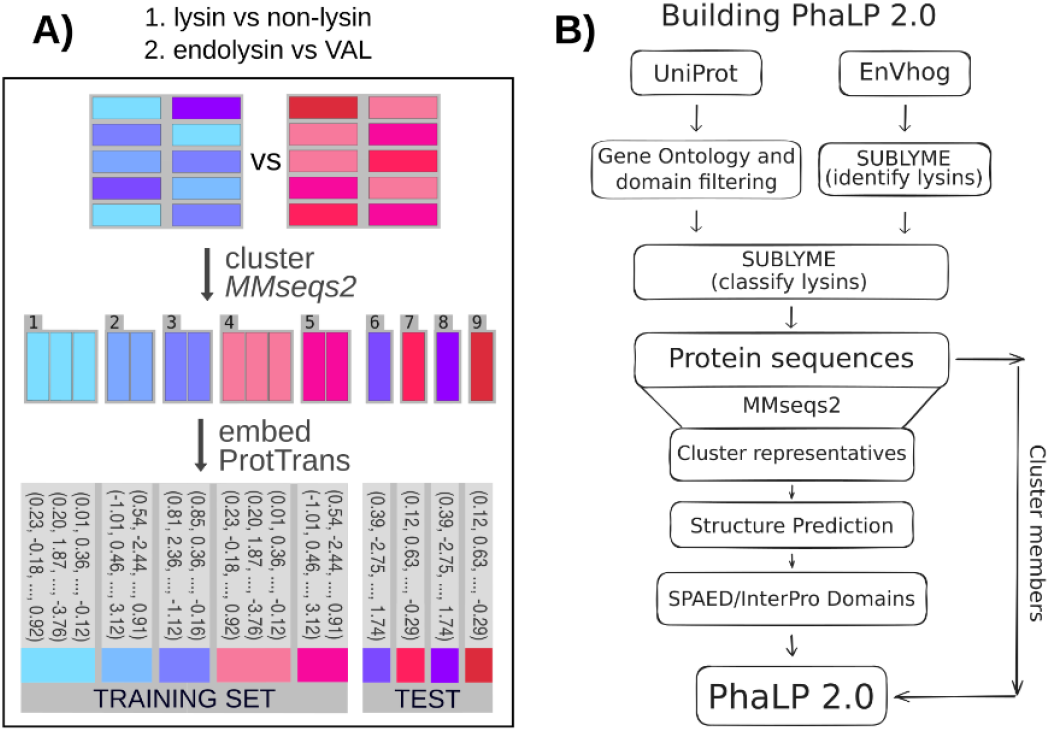
Study overview. **A)** Training SUBLYME. A first model was trained on a dataset of lysins (from PhaLP) and non-lysin phage proteins (from INPHARED) to distinguish between these two classes. Then, a second model was trained, using just the lysins in PhaLP, to classify lysins as endolysins or virion-associated lysins (VALs). Training and testing sets were split ensuring that proteins belonging to the same 30% identity (70% coverage) clusters were present in only one of these subsets, and models were trained on protein embeddings obtained from ProtT5. **B)** Schematic overview of the dataflow from public data resources to PhaLP 2.0. Lysins from UniProt are extracted using gene ontology or domain filters as described in Criel et al. (2021). Putative lysins are identified within the EnVhog dataset using SUBLYME. Both UniProt and EnVhog-derived lysins are subsequently classified as endolyins and VALs, once again using SUBLYME. The classified proteins are collected and clustered using MMseqs2 (30% identity and 70% coverage). Although all sequences are included in the database, at the moment, only the 3D structures of cluster representatives are provided along with their SPAED domain delineations and InterPro annotations.

SUBLYME consists of two binary classifiers that (1) identify lysins and (2) classify them as endolysins or VALs. Both classifiers are support vector machines (SVMs) with Radial Basis Function (RBF) kernels. Default hyperparameters were used to train all models. Performance of the first model (lysin vs non-lysin) was evaluated using repeated k-fold cross-validation with 10 repeats and 10 folds per repeat, resulting in 100 training-testing iterations. In every iteration, 90% of the data was used to train the model and the remaining 10% was used for testing. Repeated k-fold cross-validation was adopted to better evaluate the robustness of the final model, while ensuring that all proteins in the dataset were included in the evaluation. For the second model, 50 repeats with 5 folds per repeat, yielding 250 iterations, were used to evaluate the performance as fewer proteins were available: 15,387 lysins and 47,288 non-lysins were used in the first model, whereas only 4,429 VALs and 10,970 endolysins were used in the second. This sped up the training and thus made it more affordable to run more repeats, thereby improving the reliability of the performance estimates. In every case, clusters at 30% sequence identity and 70% coverage were used to separate the training and testing sets, ensuring that similar proteins (i.e., belonging to the same cluster) remained in the same subset and thus limiting overfitting as much as possible. Finally, after evaluating model robustness using cross-validation, a single final model for each classification task was trained on the entire dataset with all available labelled data, using the same architecture and default hyperparameters that had proven effective during cross-validation.

Model performance was evaluated using standard classification metrics, including F1-score, precision, and recall. These metrics were visualized as a function of the decision threshold of models, varying the latter from 0% to 100% in increments of 5%. This allowed for a detailed assessment of model behavior under different classification sensitivities, providing insight into the trade-offs between precision and recall at varying decision thresholds. Note that the default decision threshold used by the final SUBLYME models is 50%: for the lysin-prediction model for example, any prediction with a score greater than 50% is assigned as a lysin.

### Benchmarking SUBLYME

Fernández-Ruiz et al. (2018) proposed a pipeline to identify endolysins in 183k uncultured viral genomes via simple sequence similarity to a curated set of *bona fide* endolysins. The total ensemble of 3.8M proteins used in the latter work, including the 2,863 putative endolysins they identified, was retrieved to be used as a benchmark in this study. Embeddings of all 3.8M proteins were computed using ProtT5. Putative lysins were predicted by applying SUBLYME to the 3.8M proteins. All predicted lysins were clustered using MMseqs2 at 30% sequence identity and 70% coverage for further analysis. InterProScan v5.71-102.0 (Jones 2014) was used on the complete predicted lysin sequences to obtain functional annotations that were subsequently compared to known lysin-associated domains (known EADs or CBDs as described in Criel 2021 and Vázquez 2021; see domain_table.csv in our Zenodo archive).

Protein embeddings were projected and visualized in two dimensions using t-distributed stochastic neighbor embedding (t-SNE; manifold.TSNE from scikit-learn). The algorithm was applied to the full set of predicted lysins using the following parameters: n_components=2, init=‘pca’, learning_rate=‘auto’ and random_state=42. The projection was used to visualize clustering and overlap between the lysins predicted by SUBLYME and those identified by Fernández-Ruiz et al.

### Applying SUBLYME to the EnVhog database

The EnVhog database (Pérez-Bucio 2025) of metagenomically-sourced phage proteins was downloaded and used as a source of novel lysins to expand PhaLP 1.0. EnVhogDB is composed of 46M distinct proteins that were clustered into 11.9M standard sequence-level clusters (30% sequence identity and 70% coverage) and 2.2M remote homology clusters created by further clustering sequence-level clusters using the profiles (position-specific scoring matrix) generated from each cluster. The embeddings of the consensus sequences (obtained using HHBlits on the multiple sequence alignments from EnVhog clusters; Pérez-Bucio 2025) for each of these remote clusters were computed using ProtT5.

Putative lysins, endolysins and VALs were identified from the consensus sequences of the remote homology clusters using SUBLYME with a default classification threshold of 50%.

### Building PhaLP 2.0

All member proteins of the EnVhogDB remote homology clusters predicted as containing lysins as well as the UniProt-derived entries in PhaLP 1.0 (a total of 754k dereplicated sequences, 619k endolysins and 136k VALs) were included in PhaLP 2.0 (**Figure 1B**). Standard sequence-level clusters (30% sequence identity and 70% coverage) as well as remote homology clusters (as described above) were recalculated using MMseqs2 (Steinegger 2017) on all EnVhog sequences. This subsequently allowed for all UniProt-derived (PhaLP 1.0) sequences to be added using the “clusterupdate” function in MMseqs2. Note that this also allows for the cluster database to be updated in the future, without the need to recalculate all existing clusters.

For subsequent analyses, one protein (not the consensus) in each standard sequence-level cluster predicted as being an endolysin was picked by MMseqs2 as representative in order to use real sequences when running structure predictions. The 3D structures of the representatives of standard endolysin clusters were predicted using ColabFold v1.5.5 (Mirdita 2022). SPAED (Boulay 2025a), an algorithm that employs the predicted aligned error matrix from AlphaFold predictions to precisely delineate protein domains, was used with default parameters to segment and extract the domains of the representative sequences, and InterProScan v5.71-102.0 was used to annotate them. Only hits to Pfam with an E-value < 0.001 were considered for downstream analyses, but hits to other databases were also included in PhaLP 2.0.

Analyses of the composition of PhaLP 2.0 and of the organization and frequencies of domains within endolysins were performed in Jupyter Notebook and are available in our Zenodo archive. The physicochemical properties of all lysins were calculated using Biopython (1.84).

Protein structures were visualized using SwissPdb Viewer (Guex 1997).

### Database structure and features

The PhaLP 2.0 database runs on MySQL v8.0.41 using the InnoDB storage engine. It is made available on the web using Python 3.10.19 with the Django v5.2.8 and Gunicorn v23.0.0 libraries. For robustness and maintainability, MySQL and Python are each run in separate Docker containers (v28.5.1). Data querying no longer relies on BioMart as was the case in PhaLP 1.0. Instead, all querying and filtering have been fully integrated into the Django framework, enabling faster, more maintainable and more customizable operations.

## Data availability

The data underlying this article are available in Zenodo, at https://doi.org/10.5281/zenodo.17654702. PhaLP 2.0 is available at https://phalp.ugent.be.

## Code availability

Source code for SUBLYME is available on GitHub, at https://github.com/Rousseau-Team/sublyme and installable from PyPI, at https://pypi.org/project/sublyme/. All data analyses presented can be found in the notebooks available in Zenodo, at https://doi.org/10.5281/zenodo.17654702.

## Results

### SUBLYME performance on test set

SUBLYME consists of two models applied consecutively. The first identifies lysins within a set of proteins, whereas the second classifies the predicted lysins as VALs or as endolysins. The first model was trained on approximately 15k lysins and 47k non-lysins. The performance of this model was evaluated by training 100 models (10 repeats × 10 folds). The second VAL/endolysin model was trained on approximately 4.4k VALs and 11k endolysins and was evaluated using 250 models (50 repeats × 5 folds). Using the default decision threshold of 50%, average F1-scores of 97.8% were obtained for both models (95% CI [97.2% - 98.3%] and [97.4% - 98.2%], respectively). The mean precision and recall remained high (>0.85) for both models over all decision thresholds (>0 and <1) indicating the high confidence of models in their predictions (**Figure 2**).

**Figure 2.**
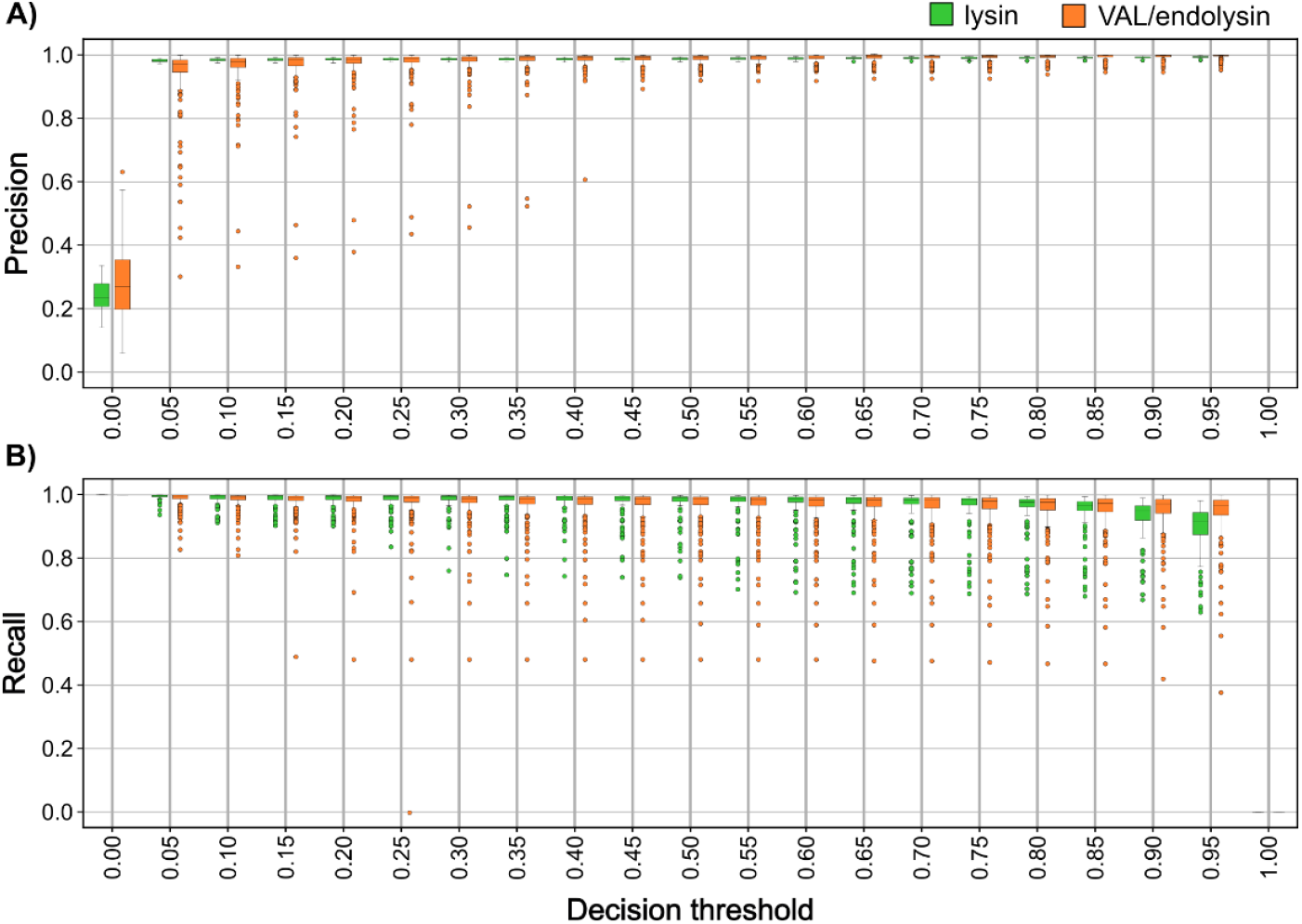
Evaluation of SUBLYME. Distribution of the precision and recall of models trained during repeated k-fold cross-validation as a function of the decision threshold used by models to make predictions. A total of 100 models (10 repeats with 10 folds per repeat) were trained and tested for the lysin prediction task (green), and 250 models (50 repeats with 5 folds per repeat) were trained and tested for the VAL/endolysin classification task (orange).

### Benchmarking SUBLYME

The Fernández-Ruiz dataset (Fernández-Ruiz 2018) was used as a benchmark to evaluate SUBLYME predictions as it had previously been used for the metagenomic mining of lysins from the 183k uncultured viral genomes (encoding 3.8M proteins) composing it. A greater diversity of lysins was identified by SUBLYME than previously reported by Fernández-Ruiz et al. (**Figure 3A**). A total of 50.8k lysins (1.3% of proteins in the dataset) corresponding to 6.8k clusters were predicted by SUBLYME, 9.8k lysins being predicted as VALs and 41k as endolysins. In comparison, 2,628 endolysins (280 clusters) had been identified by Fernández-Ruiz et al. Only 53 endolysins (∼2%) originally identified based on the homology-based approach were missed by SUBLYME.

**Figure 3.**
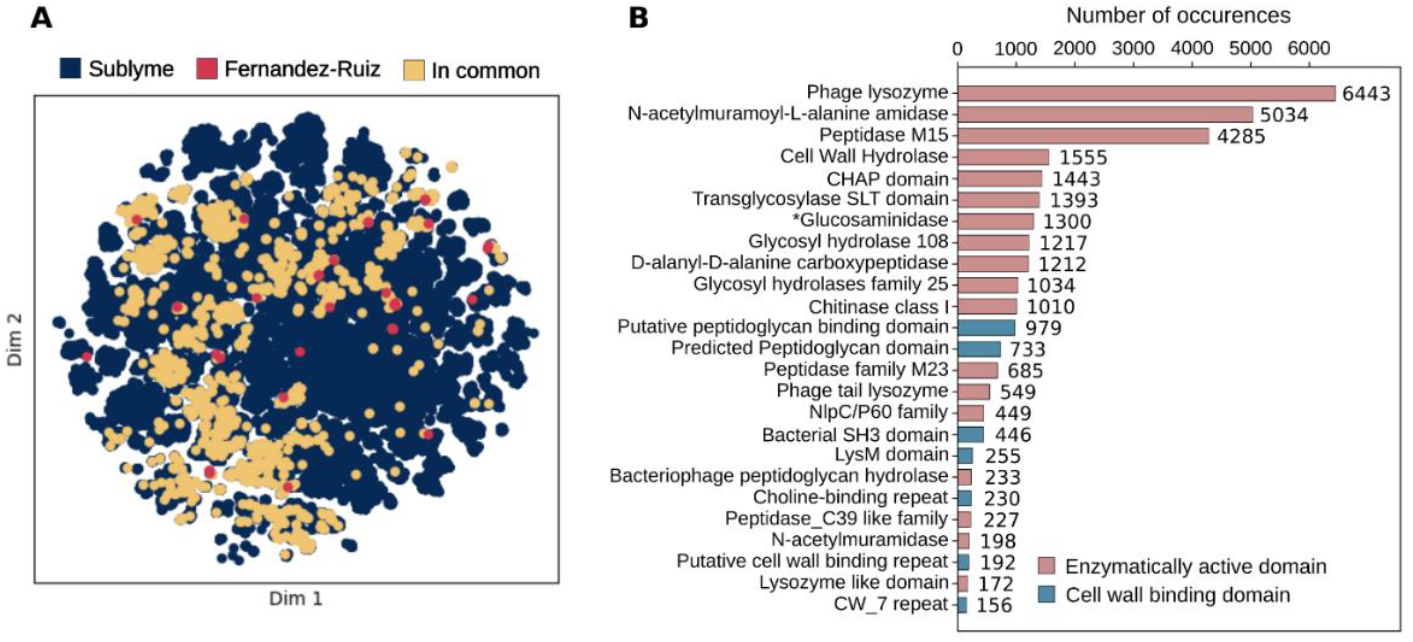
Predicted lysins in the Fernández-Ruiz dataset. **A)** t-SNE projection of protein embeddings for all predicted lysins. In blue, lysins predicted by SUBLYME; in red, endolysins identified by Fernández-Ruiz et al. (2018); in yellow, endolysins identified by both methods. **B)** Number of occurrences of the 25 most abundant domain annotations in predicted endolysins. Enzymatically active domains are in red; cell-wall-binding domains are in blue (Criel 2021; Vázquez 2021). *”Mannosyl-glycoprotein endo-β-*N*-acetylglucosaminidase” was shortened to “Glucosaminidase” as found in its corresponding Pfam entry (PF01832).

To further validate the proteins predicted by SUBLYME, they were aligned to Pfam and screened for lysin-related domains. Only endolysins were considered to simplify the analysis, since VAL domains are less known and show a much higher structural diversity. Nearly 80% (32.7k) of the 41k predicted endolysins had at least one hit to Pfam and over 98% of these were lysin-related (**Figure 3B**). This indicates that SUBLYME was able to identify a pertinent functional signal within the protein embeddings and suggests that a domain-level HMM-based approach is more sensitive at detecting lysins (32k putative lysins identified) than the sequence-homology-based approach used in Fernández-Ruiz et al. Only 135 of the predicted endolysins seemed to correspond to VALs rather than to endolysins (with domains predicted as “Type VI secretion system spike protein VgrG3”), 230 proteins had hits to domains of unknown function (DUF) and only about 150 proteins had unrelated annotations (e.g. sortase, cysteine-rich secretory protein family, target recognition domain of lytic exoenzyme, etc.). The 25 most frequent annotations are shown in **Figure 3B**.

### Building PhaLP 2.0

The EnVhog database of metagenomically sourced phage proteins was used to build the metagenomic extension of the PhaLP database. EnVhogDB is composed of phage proteins obtained from thousands of metagenomic datasets spanning diverse environments including aquatic, air, soil and host-associated samples. As such, it is the ideal resource to uncover a richer diversity of lysins. SUBLYME was applied to the consensus sequences of the 2.2M EnVhogDB remote homology clusters to identify 18.7k remote homology lysin clusters (**Figure 4A**). These correspond to 40.6k standard clusters (30% sequence identity and 70% coverage) and 743k dereplicated proteins (1.6% of all proteins in the dataset). Moreover, 621.5k of these sequences (10.6k remote homology and 25.4k standard clusters) were predicted to be endolysins, and 135.4k (8.9k remote homology and 15.9k standard clusters) were predicted to be VALs. To these sequences were added the 11k dereplicated lysins from UniProt previously contained in PhaLP 1.0, corresponding to 7.9k endolysins and 3.2k VALs. Interestingly, these sequences all clustered within the already existing clusters generated from the predicted lysin sequences in EnVhogDB. This demonstrates that the diversity in EnVhog is representative of the lysins found in UniProt and, importantly, that SUBLYME was able to correctly predict these proteins as being lysins.

**Figure 4.**
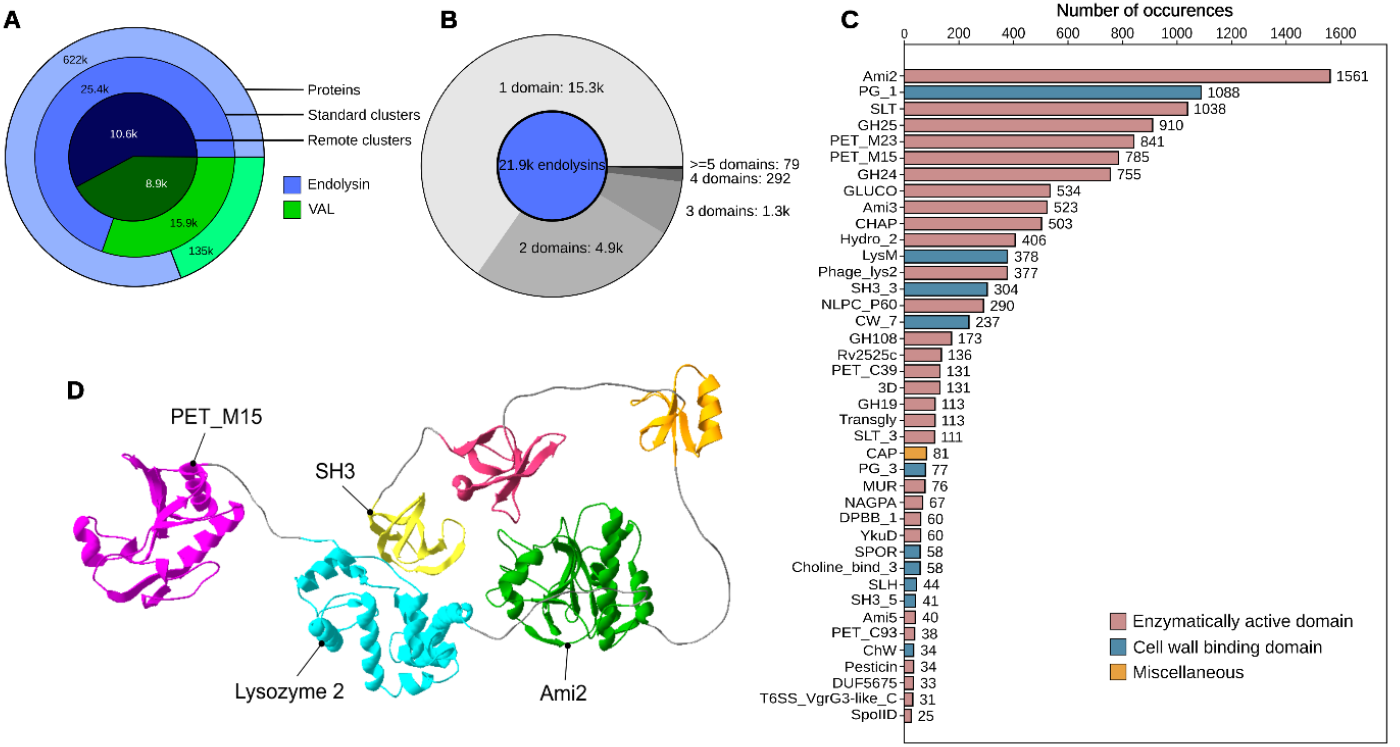
Lysins identified in the EnVhog database. **A)** The number of proteins, standard clusters (30% sequence identity) and remote homology clusters predicted by SUBLYME, split according to lysin type (endolysin and virion associated lysin – VAL). **B)** Number of representative endolysins (one per standard cluster) possessing 1, 2, 3, 4 or >=5 predicted domains. **C)** Number of occurrences of domain-level Pfam annotations identified in endolysin representatives. Enzymatically active domains (EADs) are represented in red, cell-wall-binding domains (CBDs) in blue and miscellaneous domains in yellow. **D)** Example 3D structure of a hexamodular endolysin (1G20T) possessing 3 EADs (PET_M15, Lysozyme 2 and Ami2), 1 CBD (SH3) and 2 unknown domains, visualized using SwissPdb Viewer.

For downstream analyses, one representative sequence from each standard sequence-level cluster was used to ensure a good representation of the diversity present in the dataset, while limiting the bias introduced by disproportionately abundant protein families. Using ColabFold, 21.8k 3D structures of endolysins (86%) were predicted with good confidence (pLDDT > 70), while the other 3.4k endolysin structures (14%) were removed from further analyses. Domains were identified using SPAED and subsequently compared to Pfam using InterProScan to obtain functional annotations.

Single-domain (often referred to as “globular”) endolysins are by far the most abundant class, comprising 69.5% of the dataset (15.3k clusters) (**Figure 4B**). Additionally, 22.4% (4.9k) were bimodular, 5.9% (1.3k) trimodular and 1.3% (292) tetramodular. Finally, 79 endolysins were predicted to possess five or more domains. In total, about 10.8k endolysins (49%) possessed at least one lysin-associated annotation, whereas the others contained unknown domains. Very few endolysins (<1%) solely possessed “miscellaneous” domains.

Most domains annotated by InterProScan are enzymatically active domains (**Figure 4C**), which is expected since EADs are an essential part of lysins. Ami2 (PF01510) is by far the most prevalent EAD, being found abundantly in globular as well as in modular endolysins (**Figure 4C, Figure 5**). However, the most abundant module found in globular endolysins is the SLT domain (PF01464), which is less often found in modular endolysins. GH_108 (PF05838) and PET_M15 (PF02557, PF08291, PF13539, PF05951) are abundantly found in both globular and modular endolysins and have strong associations to PG3 (PF09374) and PG1 (PF01471), respectively, unlike Ami2 which is found frequently in combination with many types of domains.

**Figure 5.**
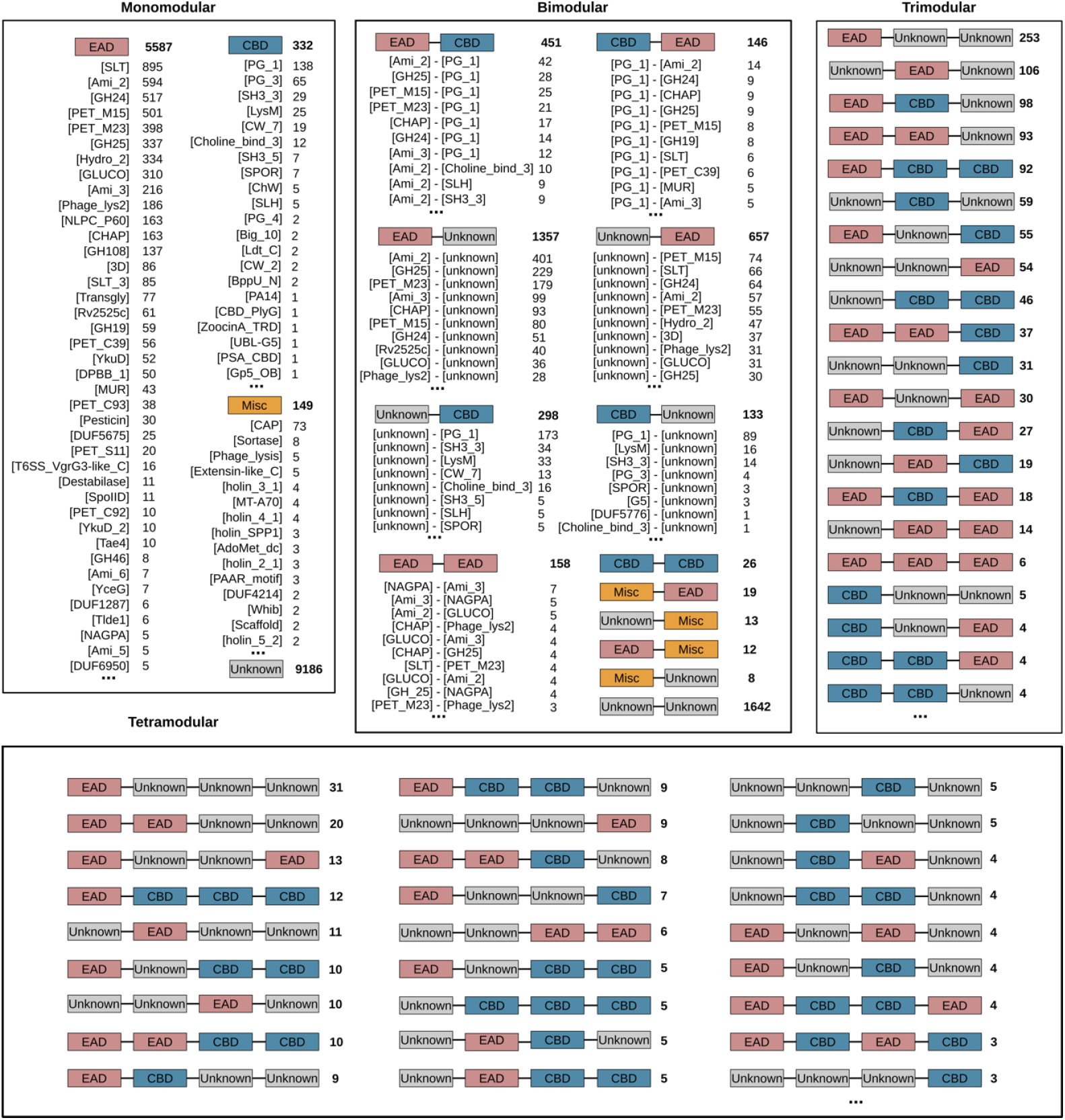
Domain architecture occurrences in PhaLP 2.0. More specific annotations are listed below the principal architecture; only the most frequent ones are listed. Enzymatically active domains (EADs) are presented in red, cell-wall-binding domains (CBDs) in blue and miscellaneous domains in yellow.

Given that endolysins must have a catalytic activity to be active, CBDs are expected to be found in modular rather than in globular endolysins. SUBLYME identified 332 globular proteins (1.5% of predicted endolysins) composed of a single CBD (**Figure 5**). These could correspond to protein fragments (Vázquez 2025) or to errors in SPAED delineations. PG1 is the most abundant CBD (**Figure 4C**). It is found in combination with nearly all the most abundant EADs (**Figure 5**) and is often found in multiple copies within the same endolysin.

Endolysins assigned a single annotation by InterProScan but predicted to have at least 2 domains by SPAED are extremely frequent in the database; 50% of predicted bimodular endolysins have an uncharacterized domain and either a known EAD or CBD (**Figure 5**). Additionally, EADs are far more often associated with uncharacterized domains (2014 occurrences, 40.9% of bimodular proteins) rather than with other annotated domains (786 occurrences, 16% of bimodular proteins). This is especially true for Ami2 which is found in 591 endolysins possessing a single annotated domain (Ami2) and at least one, but sometimes multiple, uncharacterized domains. Although CBDs are also often found solely in combination with uncharacterized domains (431 occurrences, 8.8% of bimodular proteins), they are more often found with known EADs (597 occurrences, 12.1% of bimodular proteins). In general, it is more common to find EADs in the N-terminus (40.1% of bimodular proteins have EADs in N-terminus vs 19.9% that have EADs in C-terminus), and, coherently, CBDs are more common in the C-terminus (15.8% of bimodular proteins have CBDs in C-terminus vs 6.2% that have CDBs in N-terminus).

Miscellaneous domains are relatively rare in the lysins predicted by SUBLYME, being found in just 1% of predicted endolysins. Most of these are domains of unknown function (DUF) that were frequently (99 occurrences) identified in proteins, and that could possibly be EADs or CBDs. Other domains like “sortase” and “collagen” resemble potential cell wall binding domains and “destabilase” and “holin” domains destabilize the cell wall, potentially also resembling lysin-associated domains. The presence of these domains in predicted lysins may have been caused by wrongly annotated proteins found in the training set influencing our models or by their incorporation into true lysin-containing coding sequences, perhaps due to missing stop codons.

Finally, **Figure 4D** shows an example of a hexamodular endolysin (1 of 23 found) possessing three EADs including a PET M15 domain, a lysozyme 2 domain and an Ami3 domain as well as one SH3 CBD and two unknown domains (likely CBDs judging by their structure).

### Database structure and features

PhaLP 2.0, accessible at http://phalp.ugent.be, is organized around protein clusters, enabling easy querying of cluster representatives and their members (**Figure 1B**, see also **Supplementary Figure S1** for a detailed depiction of the database). This schema allows representative proteins to store detailed information, such as domain annotations and functional data, that can be propagated to all other cluster members. In addition, this organization makes it possible to query the database for member proteins with certain characteristics using the information associated with their representative sequence.

Additionally, the “Browse database” interface has been improved with customizable table merging, offering users enhanced searchability and more intuitive access to combined datasets. A notable addition is the integration of SPAED, that enables users to explore and search for the general or specific architectures of lysins, for example searching the database for endolysins with an EAD-CBD-CBD architecture. Furthermore, the inclusion of AlphaFold structure predictions provides a more streamlined and user-friendly pipeline for protein structure analysis and can be easily exported using the “Custom table” interface.

## Discussion

PhaLP 1.0 was developed as a community-oriented database and research portal for lysin researchers. It comprised highly curated entries derived from UniProt, enriched with metadata across nine categories: proteins, phages, hosts, conserved domains, coding sequences (CDSs), gene ontologies (GOs), enzymatic activities (ECs), tertiary structures, and experimental evidence. These annotations were sourced from multiple databases, including UniProt, UniParc, NCBI Taxonomy, Virus-Host DB, InterPro, GenBank, QuickGO, ExPASy ENZYME, PDB, and PubMed. The database was designed to be highly searchable, allowing users to generate customized datasets for downstream analyses. This enabled research into novel antimicrobial molecules to be used against multidrug resistant pathogens (Yoda 2024; Paul 2025) and research into the association between lytic proteins and host range (Wang 2025) as well as facilitated new insights such as the prevalence of secondary translation initiation sites in endolysin-coding genes (Pinto 2022).

However, protein science has evolved rapidly in recent years, driven by breakthroughs in structure prediction (e.g., AlphaFold) and protein language models, alongside the exponential growth of (meta)genomic repositories, particularly for phage genomes. While PhaLP 1.0 can be updated with each new UniProt release, it remains limited to sequences from cultured phages. To address this, we developed PhaLP 2.0 (https://phalp.ugent.be), built on the foundations of PhaLP 1.0 but significantly expanded with lysin sequences derived from metagenomic sources. PhaLP 2.0 now consists of two branches (**Figure 1B**): an automatically updating stream that continues to acquire lysins from UniProt, and a newly added set of lysins predicted from the EnVhog database using SUBLYME. This second branch will also be updated when new large metagenomic datasets become available.

Incorporating metagenomic data is essential to capture the true diversity of phage lysins, especially those encoded by phages infecting unculturable hosts. Traditional homology-based annotation methods struggle with the high sequence variability found in metagenomic datasets. In contrast, embedding-based classifiers such as those used in SUBLYME leverage protein language models to extract functional signals directly from sequence data, without relying on alignment. These models outperform homology-based approaches in sensitivity and generalization (Boulay 2025b), enabling the reliable identification and classification of lysins across highly divergent sequence space. This advancement makes it possible to systematically explore lysin diversity at scale and supports the development of enriched resources like PhaLP 2.0.

In this work, we developed SUBLYME, a tool that leverages the highly informative representations provided by protein embeddings to identify lysins in metagenomic datasets. SUBLYME consists of two models: one for identifying lysins and another for classifying them as either endolysins or VALs. Both models were shown to be performant, accurately (F1-score >98%) and confidently (confidence associated to prediction >95%) classifying proteins from the testing set. SUBLYME predictions were additionally validated on an external metagenomic dataset from Fernández-Ruiz et al. Nearly all (98%) of the 2,863 previously identified endolysins in the dataset were correctly predicted by SUBLYME. Additionally, it identified 38k more endolysins (41k total) and 10k VALs, which were validated through the presence of previously described lysin-associated domains using InterProScan. SUBLYME was then applied to the EnVhog database of metagenomically-sourced phage proteins and identified 743k lysins corresponding to 40.6k standard sequence-level clusters. This represents a 40-fold increase in the number of putative lysin clusters compared to the original PhaLP database, composed of only 1,000 clusters. These newly identified lysins, along with predicted structures and domain annotations of representative sequences, were integrated to form PhaLP 2.0, a significantly enriched and scalable resource for lysin research.

While embedding-based classifiers can more sensitively assign functions than traditional alignment-based methods, they also introduce new challenges. The underlying protein language models they rely on, built on deep learning architectures, are often considered black boxes. They make accurate predictions, but their internal decision-making processes are difficult to interpret. This lack of transparency becomes particularly critical when applying such models to metagenomic datasets, which are inherently diverse and contain many sequences with no close homologs in curated databases.

One of the main challenges in this context is quantifying false positives. Models trained on lysins from cultivated phages may not generalize perfectly to the broader diversity found in environmental samples. The analyses performed here at the domain-level serve as a validation of the endolysins predicted by SUBLYME. These revealed that a substantial proportion of predicted lysins possess known lysin-related domains, specifically 80% in the Fernández-Ruiz dataset and 57% in the EnVhog dataset. This supports the reliability of SUBLYME’s predictions, while also acknowledging that some lysins lack hits to known domains, which is expected as some predicted lysins will inevitably possess very distinct sequences.

Interestingly, the proportion of endolysins with completely unannotated domains is greater in the EnVhog database than in the Fernández-Ruiz dataset, even though the proportions of proteins predicted as lysins are relatively similar in both datasets (respectively 1.6% vs 1.3%). It is important to realize that representative sequences from each cluster were used in the analysis of EnVhog, which was not the case for the analysis on the Fernández-Ruiz dataset. Selecting a single representative per cluster results in an underrepresentation of sequences from large clusters, which, incidentally, are more likely to possess previously characterized domains. The use of representative sequences from each cluster consequently increases the proportion of more distinct proteins in the analysis. Additionally, more lysins were predicted in the EnVhog database; the 40.8k sequence-level clusters are representative of a much greater diversity than the 6.8k sequence-level clusters in the Fernández-Ruiz dataset.

Most predicted endolysin clusters (69.5%) in the EnVhog database are globular. This likely reflects a greater diversity of Gram-negative associated endolysins in the dataset as globular endolysins are more typical of Gram-negative infecting phages (Oliveira 2013; Oechslin 2022). However, it may be possible that some endolysins predicted as being globular are in fact modular, but their domain boundaries and identities remain unresolved due to limitations in current annotation methods. Most endolysin clusters in the dataset are composed of one to four domains with only 130 clusters identified as having five domains or more. Among these, many exhibit repeated copies of CBDs, while others display more complex architectures. For example, the endolysin shown in **Figure 4D** possesses three distinct EADs, a CBD and two different uncharacterized domains, likely CBDs based on their position and context. Notably, no endolysins possessing two EADs of the same type were found in PhaLP 1.0 (Criel 2021). Albeit being very rare, six examples have been identified in this dataset including repetitions of PET_M23 and CHAP (identifiers: 21lwP, 5v3MF, 7DR3L, 7MKg4, T4Zi, TYor).

Domain-level observations are largely consistent with those from PhaLP 1.0, which is expected as analyses were performed by comparison to previously known domains. More importantly, many modular endolysins were found to contain at least one lysin-related domain alongside one or more uncharacterized domains. Indeed, most EADs identified in bimodular endolysins were associated with uncharacterized domains ([EAD]-[unknown]) rather than with other annotated domains ([EAD]-[CBD]), highlighting the immense functional diversity of lysins that has yet to be studied. CBDs, by contrast, are more often associated with known EADs ([EAD]-[CBD]) than with unknown domains ([unknown]-[CBD]), although the latter configuration is not rare. This asymmetry in domain annotation may reflect a gap in our understanding of CBD diversity, as many of the unknown domains paired with EADs are likely uncharacterized CBDs. This suggests that a relatively greater diversity of CBDs exists than EADs and suggests that the evolutionary space of EADs is relatively limited, at least compared to CBDs. This is consistent with the notion that evolving binding specificity is generally more accessible than evolving catalytic activity (Ulrich 1997).

To conclude, PhaLP 2.0 is the most extensive resource for the study of lysins to date. Beyond the identification and analysis of abundant and uncharacterized domain families, this database will serve in systematic comparative studies of lysin and domain evolution. SUBLYME is also available publicly for users to apply to their own datasets. Finally, amid the antimicrobial resistance crisis, these resources will hopefully also benefit the development of lysin-based therapeutics.

## Acknowledgements

AB is supported by fellowships from the FRQNT (#325947), from the CREATE Responsible Health and Healthcare Data Science (RHHDS) program from NSERC and from the Mitacs Globalink Research program. VN is funded by the Research Foundation – Flanders (FWO), grant number 1S91526N. ER is funded by a Research Scholars – Junior 1 in artificial intelligence and digital health by FRQS (#307935). RV was supported by a postdoctoral fellowship of the ‘Bijzonder Onderzoeksfonds’ (BOF), Ghent University (01P10022). This research was enabled in part by support provided by Compute Ontario (https://www.computeontario.ca/) and the Digital Research Alliance of Canada (alliancecan.ca).

## Contributions

**AB**: conceptualization (equal); software (equal); data curation (lead); writing – original draft (lead); formal analysis (equal); writing – review and editing (equal). **RV**: conceptualization (equal); formal analysis (equal); data curation (supporting); writing – review and editing (equal); supervision (lead). **VN**: writing – review and editing (equal); software (equal). **BC**: writing – review and editing (equal); software (supporting). **ER**: writing – review and editing (equal); supervision (equal). **YB**: conceptualization (supporting); writing – review and editing (equal); supervision (equal). **CG**: writing – review and editing (equal); supervision (equal). **MS**: funding acquisition; writing – review and editing (equal). **BDB**: funding acquisition; writing – review and editing (equal).

