## Supplementary material for "PhaLP 2.0: extending the community-oriented phage lysin database with a SUBLYME pipeline for metagenomic discovery"

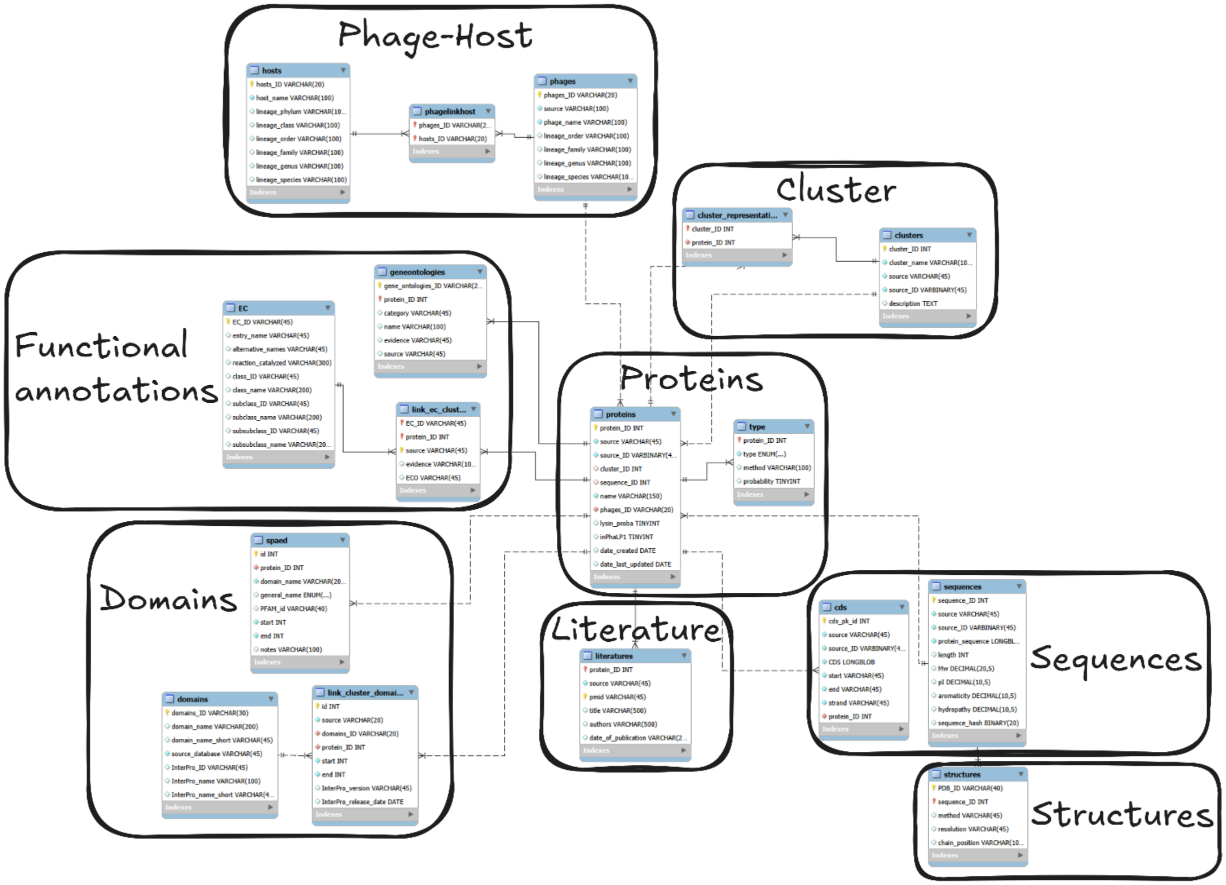


**Supplementary figure S1**. Detailed Enhanced Entity-Relationship (EER) schema of MySQL tables and relationships. The *proteins* table in the center is used to store the protein and cluster identifiers as well as the data source of each entry. The *type* table stores the lysin type of each entry ('endolysin', 'VAL'). The *clusters* table saves the cluster identifier and name, while the *cluster_representative* table connects the *cluster* table with its representative protein. The *phages* table is linked to the corresponding hosts using a *phagelinkhost* table to allow for phages to be connected to multiple hosts. The *cds* and *sequences* tables, store the Coding DNA Sequence (CDS) and amino acid sequence respectively. The *structures* table is used to link PDB identifiers to sequence identifiers. The *spaed* table stores the protein delineation together with the InterPro inferred Pfam annotations. The *link_cluster_domains* table connects the protein with the *domains* table, the latter storing other (non-Pfam) InterPro annotations. Functional annotations such as gene ontologies and Enzyme Commission (EC) classification are stored in the *geneontologies* and *EC* tables respectively. Lastly, literature data, such as PMID identifiers, are stored in the *literatures* table.
